# Cellular labelling favours unfolded proteins

**DOI:** 10.1101/274761

**Authors:** David-Paul Minde, Manasa Ramakrishna, Kathryn S. Lilley

## Abstract

Folded enzymes are essential for life, but there is limited *in vivo* information about how locally unfolded protein regions contribute to biological functions. Intrinsically Disordered Regions (IDRs) are enriched in disease-linked and multiply post-translationally modified proteins. The extent of foldability of predicted IDRs is difficult to measure due to significant technical challenges to survey *in vivo* protein conformations on a proteome-wide scale. We reasoned that IDRs should be more accessible to targeted *in vivo* biotinylation than more ordered protein regions, if they retain their flexibility *in vivo*. Indeed, we observed a positive correlation of predicted IDRs and biotinylation density across four independent large-scale proximity proteomics studies that together report >20 000 biotinylation sites. We show that biotin ‘painting’ is a promising approach to fill gaps in knowledge between static *in vitro* protein structures, *in silico* disorder predictions and *in vivo* condition-dependent subcellular plasticity using the 80S ribosome as an example.

## Introduction

Cellular complexity often arises from structurally disordered proteins^1-4^. Intrinsically disordered regions (IDRs) within proteins often overlap with sites of alternative splicing and post-translational modifications (PTMs). Both splicing and PTMs together are estimated to expand the number of proteoforms into the millions despite a relatively compact (∼20,000 large) protein-coding human genome^5-7^. Proteins rich in IDRs, Intrinsically Disordered Proteins (IDPs), are often linked to diseases such as cancer, neurodegeneration and heart diseases^8-13^. Interest in IDPs is thus increasing within the biomedical research community.

Despite increasing community interest, it has remained challenging to define the phenomenon of “intrinsic disorder” as clearly as the ordered complement of the structural proteome. Rigidly folded proteins can be solved in high-resolution crystal, cryo-EM or NMR structures that can be described by a simplified hierarchy of elements of increasing length from primary structure (sequence of single amino acid) over secondary structure elements (α-helices and β strands of ∼10 residues) to tertiary structure (folded domains of ∼100 residues) and quaternary structures (i.e. assemblies of several folded proteins). IDPs cannot be as straightforwardly classified in a simple hierarchy of modules of increasing length because the “minimal unit”, a single IDR, can vary in length from a few residues to thousands. Accordingly, IDRs can vary significantly in their properties and functions and the need for further differentiation of sub-classes of disorder was recognized early in the development of the field^14^. While the structure-function paradigm is fully established and has been highly successful, a complementary “disorder-function” paradigm is still emerging^15^.

Co-evolutionary inference suggests that many predicted disordered regions have the capacity to fold and are selected in evolution by contact constraints imposed by their folded conformation in presence of cellular binding partners^16^. In other words, such binding-coupled folding IDPs look similar to folded proteins as determined by (co)evolution statistical analysis. Interfaces of foldable IDRs tend to be larger than contacts between two ordered proteins and the exposed hydrophobic surface area is often larger, which in some cases limits solubility of IDPs and requires tighter subcellular regulation of IDPs compared to ordered proteins^17-19^.

One of the least characterized aspects in IDR research is *in vivo* malleability leading to multiple structural forms that disordered regions can adopt in a given compartment in a given cellular state. According to *in vitro* experiments, it can be expected that subtle variations in pH, salt concentrations, and PTMs can have very significant effects on the conformational ensembles of IDPs. For instance, nuclear pore proteins can form extremely tight complexes (dissociation constant (K_d_) in low pM range) near physiological salt concentration (∼100 mM) which becomes very weak (K_d_ in mM range) at 200 mM salt concentration^20^. Indeed, a recent large-scale multidimensional proteomics study that investigated temperature-dependent solubility and abundance changes across cell cycle phases, demonstrated that large subsets of the human proteome dramatically change their solubility, stability, subcellular organization and protein partners in patterns resembling differential phosphorylation during the cell cycle^21^.

Early reports suggested that phosphorylation predictions can become significantly more accurate if local intrinsic disorder tendency is taken into consideration^22^. Many single-protein examples illustrate that IDRs can be phosphorylated or hyper-phosphorylated within disordered residues, often at highly soluble and intrinsically disorder-promoting serine and threonine residues^10,11,23-25^. The correlations of IDRs with acetylation, ubiquitination and sumoylations at lysines, and phosphorylations at residues such as tyrosine and histidine are more challenging to detect, however, and hence frequently underreported in scientific studies^26-30^. Finally, there are very few studies reporting possible interactions between IDRs and multiple types of PTMs.

Biotinylation-based proximity proteomics methods are traditionally used to map transient interactions and subcellular neighbours^31-35^. The common principle of various proximity proteomics approaches is that biotinylation is highest in proximity to the biotin-activating enzyme that is fused to protein of interest. This localised biotinylation enhances the biotin incorporation in protein interactors and/or subcellular neighbours of biotin activating enzyme fused targets, which can be quantified in mass spectrometric experiments combined with stringent statistical filtering of background proteins to remove endogenously and non-specifically biotinylated proteins. Several recent technological improvements enable the direct detection of thousands of biotin sites in hundreds of proteins in a single study^36-39^. We therefore reasoned that these novel large-scale *in vivo* biotin site data could be repurposed to gain insights into possible cellular conformations of proteins.

The most frequently used enzymes in proximity proteomics are variants of BirA biotin-protein ligase and Ascorbate Peroxidase (APX)^33,34,40,41^. A promiscuous mutant of BirA (“BioID”) as well as a thermophilic homologue (BioID2) biotinylate nearby lysines through the formation of activated biotinoyl-5’-AMP which forms a covalent attachment to the nucleophilic ε-amino side chain group of lysine (K). APX or accelerated versions like APEX2 can convert biotin-phenol to activated radicals that can readily react for a short period of time with nearby tyrosines (Y). Interestingly, these two amino acid types are on opposite ends of the disorder-promoting amino acid scale — Lysine promotes disorder while tyrosine is on average depleted in IDRs^42^.

We hypothesize that sites of cellular biotinylations in proximity labelling studies could favour biotinylation within predicted IDRs if these can retain greater accessibility in their cellular context compared to predicted ordered regions. We perform a comprehensive analysis using representative data from multiple, independent and orthogonal large-scale proximity tagging datasets as well as diverse IDR predictions to test our hypothesis. We demonstrate the enrichment of cellular biotinylation events in predicted IDRs and show that these regions show higher biochemical reactivity compared to ordered regions in all targeted cellular niches and especially in the nucleus of HEK293 cells.

## Results

### Concept of the study

Predicted IDRs can be reshaped by interactions in cells (Fig. 1A). Often, specific functions of IDRs are linked to their potential to fold upon interacting with specific native partner proteins or ligands ^43^. Alternatively, short “linear motifs” within IDRs could mediate a multitude of local interactions of other folded proteins that could constrain and compact IDRs in specific partially folded or ordered conformations^44^. In the most extreme scenario, IDRs could remain entirely unfolded and fully accessible^20,45^. We expected to observe more *in vivo* biotinylations within predicted IDRs if they remain at least transiently and locally unfolded and accessible in their cellular context. If present, such a correlation can be used for *in vivo* structural proteomics studies.

**Fig. 1.**
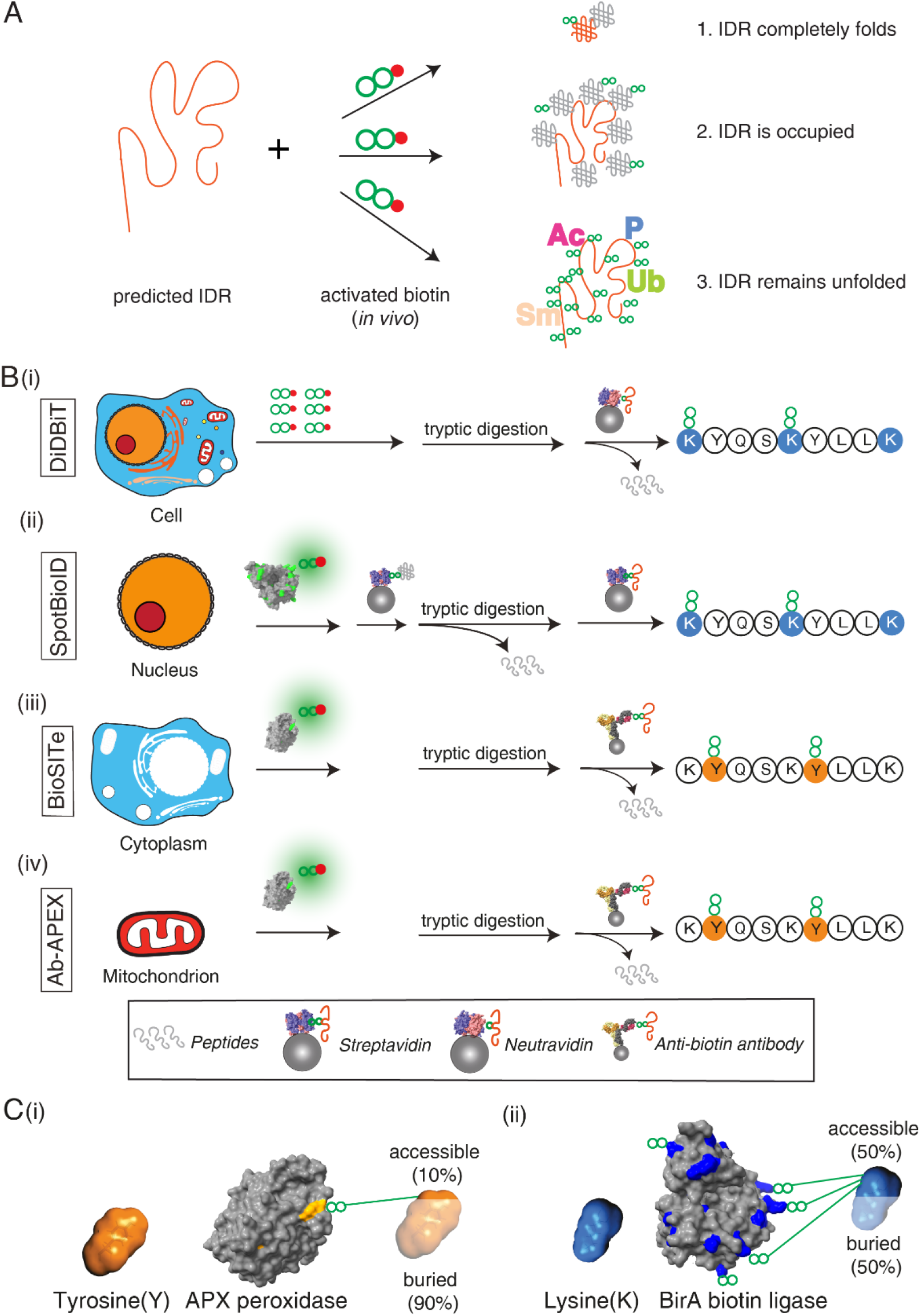
Large-scale *in vivo* biotinylation datasets can be re-purposed to identify accessible protein regions *in vivo*. (**A)** Concept of the study. Predicted IDRs are compared with *in vivo* biotinylation sites and the most frequently reported post-translational modification sites to identify highly accessible regions in proteins. (**B**) Study design of re-analyzed studies (i) DiDBiT^38^ (ii) SpotBioID^39^ (iii) BioSITe^36^ (iv) AB- APEX2^37^. (**C)** (i) Tyrosine solvent accessible surface area (SASA) is reduced significantly (∼90%^47^) in folded proteins as illustrated on the APX structure (PDB ID 1APX) (ii) Lysine sidechains contribute a large fraction of the total surface in typical folded proteins as illustrated in the *Aquifex aeolicus* birA (BioID2) structure (PDB ID 3EFR). Some 50% of the average lysine’s SASA stays exposed in folded proteins^47^.

### Brief introduction to selected proximity proteomics studies

To test our hypothesis of possible links between structural features of proteins and biotinylation, we selected four recent, independent and orthogonal, large-scale studies by the following criteria (1) large number of directly identified biotinylation sites (2) orthogonality in targeted subcellular niches and (3) independence of biotin-peptide enrichment strategies (Fig. 1B). “DiDBiT” targeted the whole cell and is therefore agnostic of subcellular localisation. It identified ∼20 000 biotinylation sites on lysine sites upon extensive biotinylation by applying 1mM NHS-biotin, a chemically activated form of biotin, to cultured HEK293 cells, complete digestion by trypsin and streptavidin-affinity purification of biotinylated peptides^38^. “SpotBioID” targeted rapamycin-dependent interactions of the human mTOR kinase using its FK506-rapamycin binding (FRB) domain fused to BioID^39^. Immunofluorescence data within SpotBioID and previous literature conflict concerning the main subcellular localisation of FRB- BioID that appears to be cytoplasmic in fluorescence experiments and nuclear in previous literature and biotin-protein enrichments^39^, with most evidence suggesting mainly nuclear localisation of the FRB-BioID fusion. The remaining two data sets come from recent, tyrosine-targeting APEX2 studies. Both successfully explored an alternative enrichment strategy based on polyclonal biotin-antibody from goat and rabbit that facilitated gentle elution while retaining explicit biotin site information unlike other strategies involving gentle elution of cleavable biotin derivatives^36,37,46^. They comprise an antibody-based APEX2 study (within this paper termed “Ab-APEX”) targeted the mitochondrial matrix using mito-APEX2^37^, and a study called BioSITe^36^ which uses a cytoplasmic APEX2 fusion construct to Nestin (NES) protein.

### Orthogonality of tyrosine and lysine as molecular targets of proximity proteomics

How different are tyrosine and lysine residues, the most frequent molecular targets in proximity proteomics? Tyrosine is a partly hydrophobic and bulky amino acid and predominantly partitions to the hydrophobic core of proteins and near the interface of intrinsic membrane proteins. Its solvent accessible surface area (SASA) shrinks by some 90% during folding reactions (Fig. 1Ci)^47^. Lysine residues, by contrast, tend to orient to the surface of folded proteins and stay in contact with surrounding water molecules, i.e. retain a large fraction of their SASA (Fig. 1Cii). Nevertheless, through their intramolecular and intermolecular contacts, for instance, in protein-protein interactions, lysine residues have a large spectrum of accessibilities with an average near 50% of remaining SASA in folded proteins^47^.

### Proximity proteomics studies can specifically target subcellular locations

As expected, as the four studies targeted different subcellular niches there was a very small overlap in proteins across the 4 studies with only 29 proteins being in common (Fig. S1A). Of these 29, many of them had multiple cellular locations predominating in the nucleus (Fig. S1B, blue), cytosol (Fig. S1B, red) and the extracellular region (Fig. S1B, yellow). Given the small size of this subset of the whole dataset, these locations are not statistically enriched for despite being frequently seen. However, we could confirm the location for each of the studies above (n > 500) using a functional enrichment analysis against a set of Gene Ontology (GO) terms aimed at describing cellular location (GO:CC; Fig. S1C). Our data shows that as expected, Ab-APEX proteins strongly target the mitochondrion with high fold enrichment for the mitochondrial matrix and the mitochondrial inner membrane (Fig. S1C, first column). Then, we checked the BioSITe data which also as expected based on the NES-APEX fusion, enriched GO terms of the cytoplasm and the cytosol (Fig. S1C, second column). The DiDBiT study, which lacks specific targets seems enriched for nuclear, mitochondrial and cytosolic proteins (Fig. S1C, column 3). Finally, SpotBioID, where the authors state that FRB-BioID is cytoplasmic, are enriched for mostly nuclear and some cytoskeletal proteins^39^. Briefly, all four studies showed expected enrichments consistent with the targeted cellular compartment and previous literature.

### Illustrative examples of proteins that are biotinylated across all four independent studies

We started by exploring our datasets combining all biotinylation sites and PTMs by initially focusing on structural features of the limited subset of 29 proteins that were common in all studies. While not statistically significant, we noticed that the list contained many RNA binding proteins. Elevated IDR content among these proteins is consistent with previous reports of high IDR content among nucleotide-binding proteins^48^ but a larger set will have to be explored for firmly establish a statistical correlation. The first example, Emerin, is an integral membrane protein that is often found at the inner nuclear membrane or at adherens junctions. Emerin mutations cause X-linked recessive Emery– Dreifuss muscular dystrophy. Biotinylation sites from all four studies cluster in a large predicted IDR in the first half of the protein sequence, avoid the transmembrane-spanning domain (Fig. 2Ai-iv) consistent with our hypothesis that predicted IDRs might are more biotinylated *in vivo* if they remain highly accessible. A very large number of other PTMs in this IDR further illustrates that this membrane protein is indeed often subjected to intracellular modifications. Surprisingly, Emerin is found in all four studies despite the fact that some targeted different subcellular locations. Emerin is one of ∼400 identified integral membrane proteins, suggesting that detailed intracellular structural insights can be gleaned from re-purposed proximity proteomics studies.

**Fig. 2.**
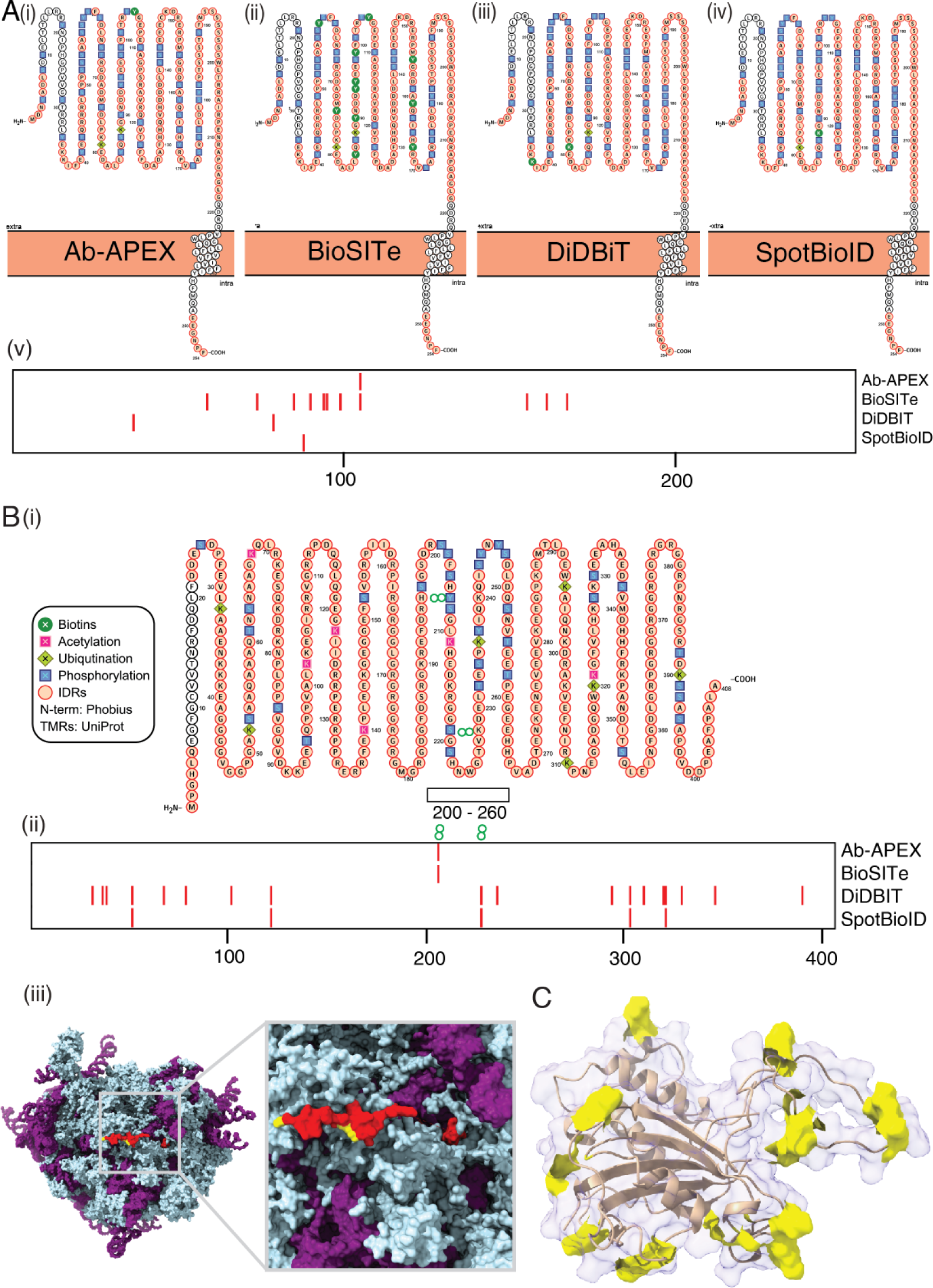
Illustrative examples for *in vivo* surface biotinylation is four independent studies. (**A)** Emerin (i-iv) Protter representations of regions of IDR (orange), frequent PTMs and biotin modification in the four studies (v) comparison of biotinylation sites across four studies. (**B)** SERBP1/Plasminogen activator inhibitor: (i) Protter plot showing post-translational modifications and regions of IDR (orange) (ii) sites of biotinylation across four studies (iii) Cryo-EM model (PDB ID 4V6X^70^) of SERBP1 with sites of biotinylation across all studies highlighted in yellow and the visible fraction (∼20%, most coil and α- helix) of the Plasminogen activator inhibitor 1 RNA-binding protein highlighted in red (**C)** FKPB3 NMR structure (PDB ID 2mph) with biotinylations in yellow, non-biotinylated chains are represented in cartoon style under 90% transparent surface.

Next, we analysed the predicted fully disordered RNA-interacting plasminogen activator inhibitor protein SERBP1 (Fig. 2Bi). Four sites of biotinylation, across the four studies, cluster around the central region of this protein (residues 200-260) where previously reported unique PTM sites also cluster (Fig. 2Bii). DiDBiT identifies many additional sites scattered over the entire protein sequence, five of which are common with the nuclear targeted SpotBioID study. SERBP1 was previously found in multiple subcellular locations consistent with its identification in four studies. Ribosomal proteins are typically predicted to be disordered or non-globular^49^. We see SERBP1 attaching at the periphery of the 80S ribosome RNA-protein complex and mostly lack (in ∼80% of its sequence) unique electron density (Fig. 2Biii); remaining small visible fractions form elongated structures that are detected in random coil or α-helical conformations.

Finally, we selected FKBP3 as a protein of average (predicted) disorder content for the human proteome around 40% according to VSL2b^2^. FKBP3 is a cis-trans prolyl isomerase that is involved in cellular protein folding and tightly binds to the immunosuppressant rapamycin. Biotinylations are enriched in its predicted IDRs (72%) or localise to local coil structure and short, highly accessible α- helical segments in the NMR structure. FKBP3 was previously annotated as nuclear protein. We conclude that detailed inspection of common examples across four studies suggests an enrichment of biotinylations in IDRs and regions lacking defined secondary structure in otherwise folded proteins.

### Predicted IDRs are more frequently and densely biotinylated in vivo

Encouraged by observing enhanced biotinylation in predicted IDRs in the small pool of proteins common to all four studies, we next wondered whether this trend might still hold globally for the biotinylated proteome (referred to as “biotinome” hereafter) comprising nearly 4000 proteins. We first checked if proteins with higher predicted fraction of IDRs contain higher numbers of unique sites of biotinylation by comparing the predicted IDR fraction for proteins in each biotinome to the number of biotinylated sites they contained (Fig. 3A). Within each dataset, there were only a small number of proteins with 5 or more biotinylation sites and hence these have been collectively binned into the “5+” category (Fig. 3A, last violin). For both the SpotBioID and the BioSITe studies, we observed an increase in the frequency of sites of *in vivo* biotinylations per protein from 1 to 4 with increasing IDR fractions, while DiDBiT and Ab-APEX did not show this trend (Fig. 3A, left panels). Both the cytosol and the nucleus, which are target compartments in BioSITe and SpotBioID have been previously suggested to contain many IDRs^48^. Mitochondria, by contrast, are predicted low in IDRs especially their subset of proteins with bacterial homologues^50^. DiDBiT, lacking compartmental preference, contains both highly disordered and fully folded proteins which might mask any possible weak correlation. We conclude that *in vivo* observed biotinylation frequency per protein and predicted IDR fractions can be correlated in IDR-rich compartments such as the nucleus and cytosol in HEK293 cells (Suppl. Table “Biotins”).

**Fig. 3:**
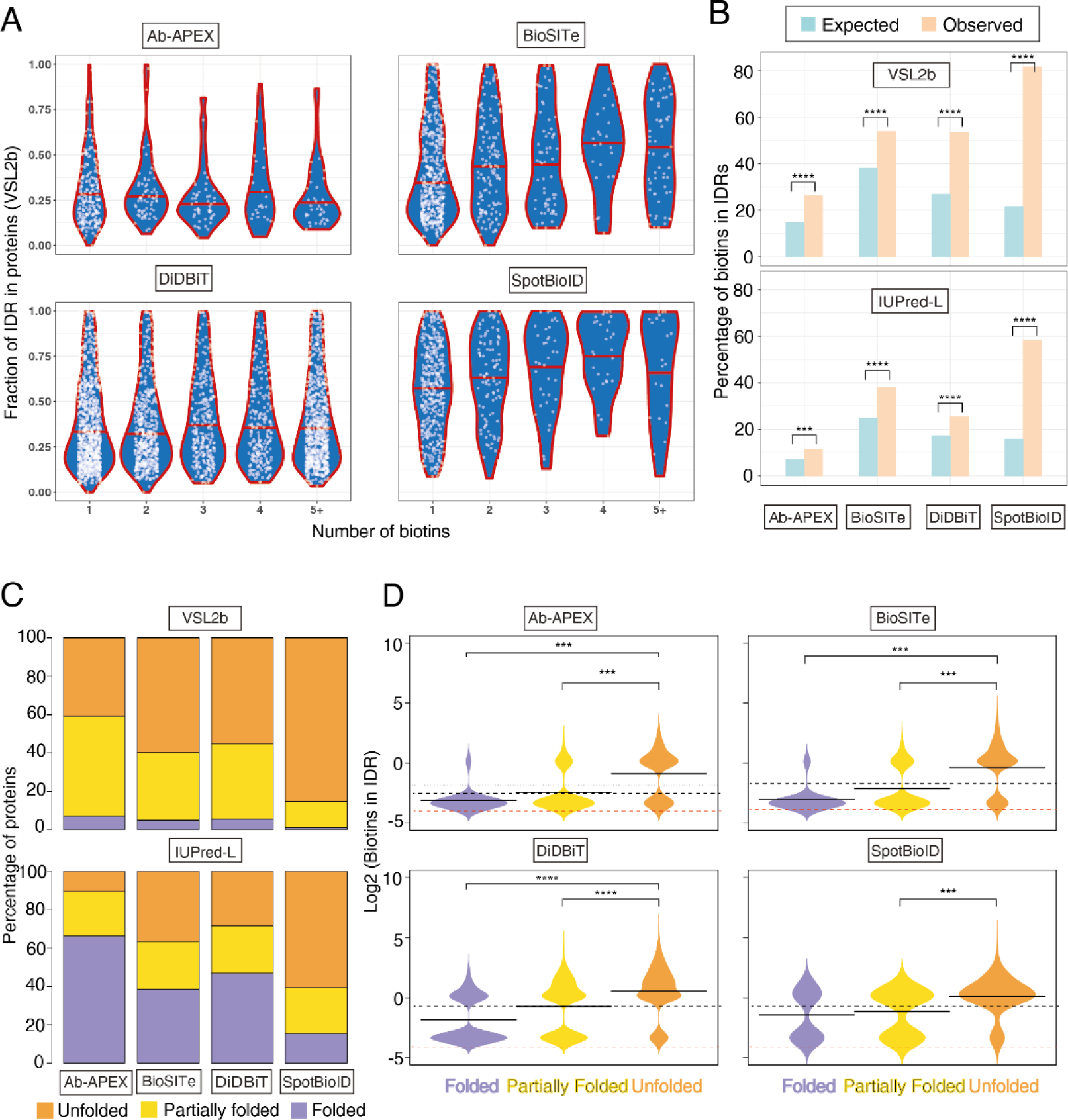
Predicted IDRs are preferentially *in vivo* biotinylated across all studies. **(A)** Violin plots (i.e. mirrored density distribution plots) showing the relationship between number of biotins and predicted fraction of IDR across all biotinylated proteins. Biotin numbers >=5 are grouped into one set to show the general trend of the data. The red line in the middle of each violin represents the median fraction of IDR for that group. **(B)** Barplots showing the Expected (pale blue) and Observed (pale orange) distribution of biotins within regions of IDR across the 4 studies using the IDR caller VSL2b (Top) and IUPred-L (bottom). Significance: ***p < 0.0005; ****p close to 0, using a bionomial test where the “probability of success” is the (number of lysine residues or tyrosine residues in IDRs/Total number of lysine residues or tyrosine residues), a “success” is a biotin within an IDR and “number of trials” is the number of biotins observed in that study. **(C)** Barplots showing the distribution of proteins from the 4 studies across the 3 structural classes^18^: Folded (F, 0-10% disorder; purple); Partially Folded (P, 10-30% disorder; Yellow) and Unfolded (U, > 30% disorder; orange) for two different IDR callers VSL2b (Top) and IUPRed-L (bottom). The numbers of proteins in the VSL2b caller are displayed in S2B **(D)** Bean-plots^72^ showing the distribution of biotins that occur within IDRs across the 3 classes F, P, U in each study. The y-axis in on a log2 scale with values 0 and above representing 1 or more biotins. The red dotted line represents 0 biotins on a log scale (with added correction factor). The solid black line in the middle of each violin represents the mean biotins (on log2 scale) for that group. The black dotted line represents the mean log2(Biotins) across all groups.

To overcome limitations of averaging over IDRs and ordered regions that might have masked structural trends in the DiDBiT and Ab-APEX studies, we next refined our analysis by distinguishing between biotinylations inside and outside of IDRs while accounting for the density of potentially modifiable residues. To establish an “expected” rate of biotins, we calculated the number of lysine residues (K; for SpotBioID and DiDBiT) or tyrosine residues (Y; Ab-APEX and BioSITe) - both within the predicted regions of IDR (as determined by VSL2b) and across the entire protein body. The ratio of all K/Y residues within IDR regions to all K/Y residues across the protein body gave us an expected rate of biotinylation in IDRs. We then performed a similar calculation using the numbers of biotins we actually observed within IDRs and across the whole protein for each of our 4 studies (Fig. 3B). Consistent across all 4 studies, irrespective of the prediction algorithms used, we observed a significantly greater number of biotins within IDR regions (orange bars; Fig. 3B) than expected (blue bars; Fig. 3B). Once again, this observation was more significant in the nuclear proteins (SpotBioID) than in the mitochondrial proteins (Ab-APEX) (Fig. S2A).

Convinced that we are seeing a true positive correlation between local predicted IDRs and biotinylation density, we sought to see if similar trends can also be observed on protein level after sorting all proteins in classes ranging from most to least folded. To this end, we labelled a protein as Folded (F) if it had predictions of <10% IDR, Partially Folded (P) if it had 10-30% IDR and Unfolded (U) if it had >30% IDR in its protein body similar to a strategy in Gsponer et. al.^18^. We then looked at the overall distribution of proteins in these IDR classes for each of our 4 studies (Fig. 3C). We display the results for just VSL2b and IUPred-L algorithms as “VSL2b_IUPred-L” mimics the trend of VSL2b alone while the “D2P2 consensus” mimics IUPred-L. We observed that all studies contain proteins that can be classified as F, P and U thus enabling pairwise comparisons. The predictors that are better at predicting long IDRs or the absence of folded domains, IUPred-L and D2P2 consensus predictors^51,52^, classified more proteins as F than VSL2b that has a wider definition of IDRs that also includes short IDRs. Consistent with our previous observations and claims in literature^20,48^, the nuclear protein enriched SpotBioID dataset shows the highest proportion of U proteins while the mitochondria targeting Ab-APEX study shows the highest proportion of F proteins (Fig. 3C, S2B).

Given these three categories of proteins, we wondered whether there would be an association between IDR-associated-biotins and the various categories of IDPs. To assess this, we performed both pairwise t-tests between the groups (F-P, U-P, U-F; S2Ci) and an ANOVA across all groups followed by a Tukey’s Honestly Significant Differences post-hoc test (Fig. S2Cii). In all studies except SpotBioID, there were significant differences between biotin numbers in the F and U group with more biotinylation events occurring in the U group. Additionally, the differences were significant for all studies between U and P groups, once again showing higher number of biotins in the U group (Fig. 3D; S2C).

### Post Translational Modifications (PTMs) enriched in biotinome-IDRs

Having discovered a strong correlation between IDRs and increased biotinylation, we wondered whether an up-to-date comparison with ∼305,000 PTMs in PhosphoSitePlus comprising the small phosphorylation, acetylation as well as the larger protein-sized sumoylations and ubiquitinations, would parallel these trends or show enrichment in other proteins that do not overlap with the biotinome. To this end we downloaded all experimentally reported phosphorylation, acetylation, ubiquitination and sumoylation data from PhosphoSitePlus and mapped them to two datasets (1) the “biotinome” for HEK293 which is the collection of all proteins across our 4 studies and (2) the HEK293 proteome which was published by Geiger et. al. in 2012 and contains 7650 proteins^53^. Additionally, we also mapped IDRs to the Geiger et. al. proteome using the VSL2b algorithm.

As a simple starting point, we looked at the direct correlation between the number of IDRs and the number of PTMs in both datasets (Fig. 4A, Suppl. Table “PTM list”). In both cases, there is a modest positive correlation of 0.37 which was supported by a highly significant p-value (p = 2.2e-16) indicating that the probability of seeing this correlation by chance is extremely low. We thus conclude that there is a small but significant correlation between IDRs and PTMs in the overall proteome as well as the “biotinome”.

**Fig. 4:**
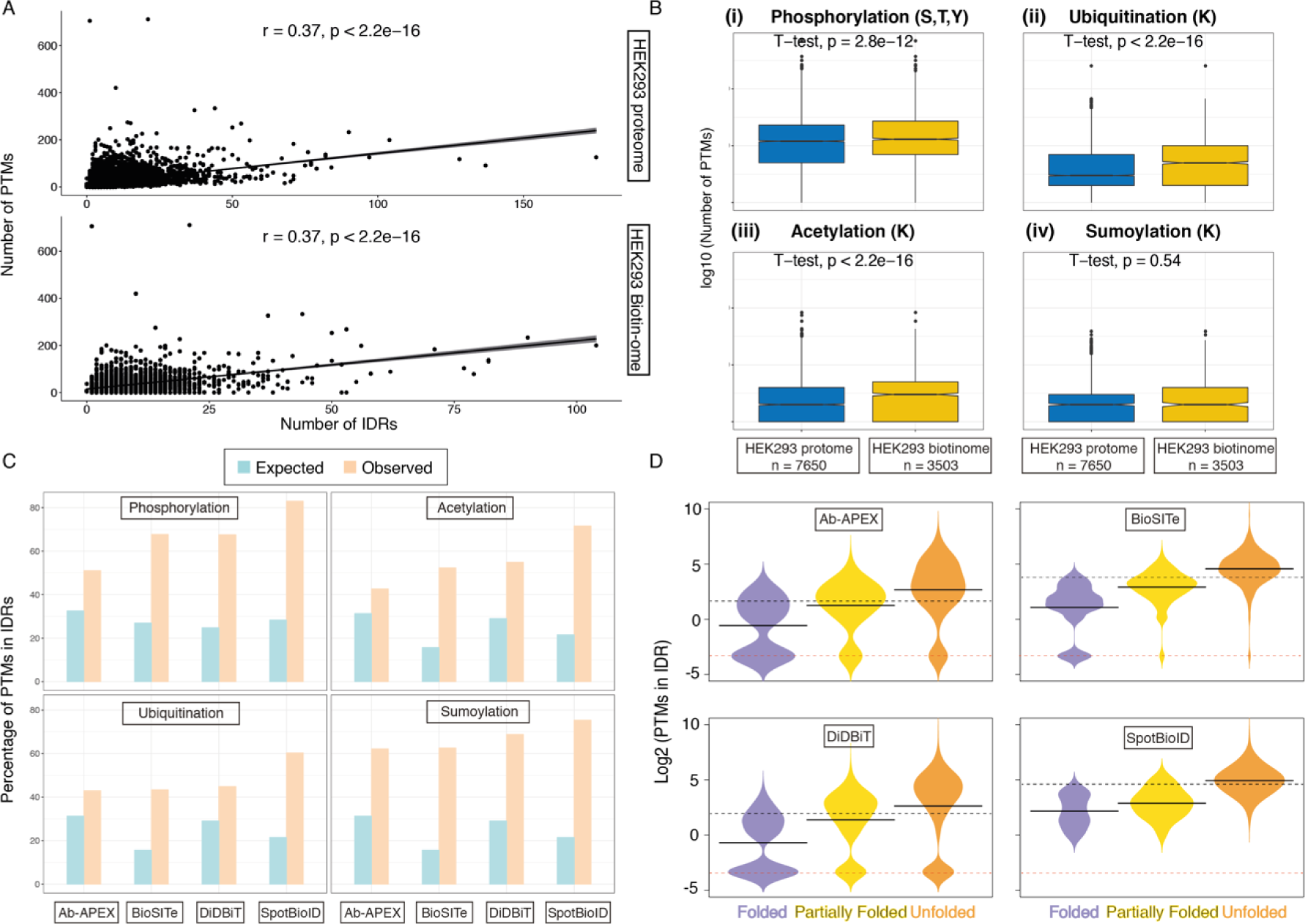
PTMs more enriched in biotinome-IDRs. **(A)** An x-y plot showing the Number of IDRs (x-axis) vs Number of PTMs (y-axis) for a published HEK293 proteome containing 7650 proteins (top panel) and for all proteins across the 4 studies (bottom panel; n = 3503). A Pearson’s correlation co-efficient is included in the along with a confidence p-value. Both relationships are positive and highly significant. **(B)** Boxplots showing the number of PTMs in the known HEK293 proteome (blue boxes) versus the “biotinylated” HEK293 proteome captured in the four datasets used in this study (yellow boxes). The four panels represent the four different post-translational marks – phosphorylation (top left), ubiquitination (top right), acetylation (bottom left) and sumoylation (bottom right). A t-test for differences in means between the two groups was conducted and the p-value is embedded at the top of each panel. All except sumoylation are significantly different and greater in the HEK293 “Biotinome” relative to the whole proteome. **(C)** Barplots showing the Expected (pale blue) and Observed (pale orange) distribution of PTMs within regions of IDR across the 4 post-translational marks using the IDR caller VSL2b. All pairs except are significantly different between Observed and Expected values using a bionomial test where the “probability of success” is the (number of K/S/T/Y in IDRs/Total number of K/S/T/Y), a “success” is a PTM within an IDR and “number of trials” is the number of (each type of) PTMs observed in that study (Fig. S3A) **(D)** Bean-plots^72^ showing the distribution of PTMs that occur within IDRs across the 3 classes F, P, U in each study. The y-axis in on a log2 scale with values 0 and above representing 1 or more PTMs. The red dotted line represents 0 PTMs on a log scale (with added correction factor). The solid black line in the middle of each violin represents the mean PTMs (on log2 scale) for that group. The black dotted line represents the mean log2(PTMs) across all groups.

Despite the similarity in correlation, we wanted to know if there was an overall enrichment of PTMs in the HEK293 biotinome relative to the HEK293 proteome. We looked for a difference in the mean number of PTMs in the two groups of proteins, across each of the 4 post-translational marks. Median frequencies followed the expected higher rates for frequently reported phosphorylations and less frequently studied and likely under-published ubiquitinations, acetylations, and sumolyations. (Fig. 4B). Furthermore, on average, there are significantly more phosophorylation, acetylation and ubiquitination marks in the HEK293 biotinome relative to the HEK293 proteome (p << 0.05; Fig. 4B, i-iii) suggesting a trend in co-occurrence of PTMs with biotins. This trend was not observed for sumoylation possibly due to very low reported numbers and technical under-detection of this specific modification (Fig. 4B, iv).

Having observed that we have a positive correlation between IDRs and PTMs, and that PTMs are more frequent in the biotinome, we wanted to see if there was a preponderance of biotinome PTMs within regions of disorder (IDRs). We calculated the expected rate of seeing serine (S), threonine (T) and tyrosine(Y) in IDRs vs the rest of the protein to work out the baseline probability of phosphorylation marks. Similarly, we calculated the expected rate for lysine in IDRs vs the rest of the protein to establish a baseline for ubiquitination, acetylation and sumoylation marks. We then calculated the observed frequency for all 4 PTM types within IDRs in our biotinome datasets. Our observations were even more remarkable than those seen in the biotin context with all 4 marks being significantly enhanced within regions of intrinsic disorder more than expected (IDRs predicted by VSL2b; Fig. 4C, Fig. S3A).

Knowing that there was a significant enhancement of PTMs in regions of IDR, we sought to determine if this would be even stronger if we looked at the proteins in the 3 previously discussed categories of Folded (F), Partially Folded (P) and Unfolded (U). We confirmed that in all 4 studies, PTMs occurred at a significantly higher rate in U proteins than in F or P proteins (Fig. 4D, S3B, S3C). This analysis also showed that all proteins in the nuclear SpotBioID study (Fig. 4D (ii)) and most of the proteins in the cytoplasmic BioSITe study (Fig. 4D (iv)) contain one or more PTMs (y-axis > −3) while this was not true for the Ab-APEX and DiDBIT studies. Given our previous observations that IDRs are more frequent in cytosol and nucleus, this provides another line of evidence that PTMs, like biotins, prefer IDR rich proteins.

### Application of biotin ‘painting’ to investigate the in vivo plasticity of the 80S ribosome

To investigate whether large structured complexes can also be analysed with this method, we filtered the DiDBiT dataset for ribosomal proteins and visualised all biotinylated subunits in an “exploded” version of the 80S ribosome (Fig. 5). Virtually all biotinylated subunits are non-spherical and multiply biotinylated as evident from large fractions of yellow marked biotinylation sites, many of which are inaccessible to water or larger molecules such as biotin in the fully assembled 80S ribosomal complex as they are contacting ribosomal RNA (supplementary video). We observed a high density of biotinylation sites in this ∼3 megadalton large complex which suggests that biotin ‘painting’ has no fundamental size limitation. High biotinylation density in the 80S ribosome is consistent with an earlier suggestion that eukaryotic ribosomes are rich in predicted IDRs that can be functionally essential^49^.

**Fig. 5.**
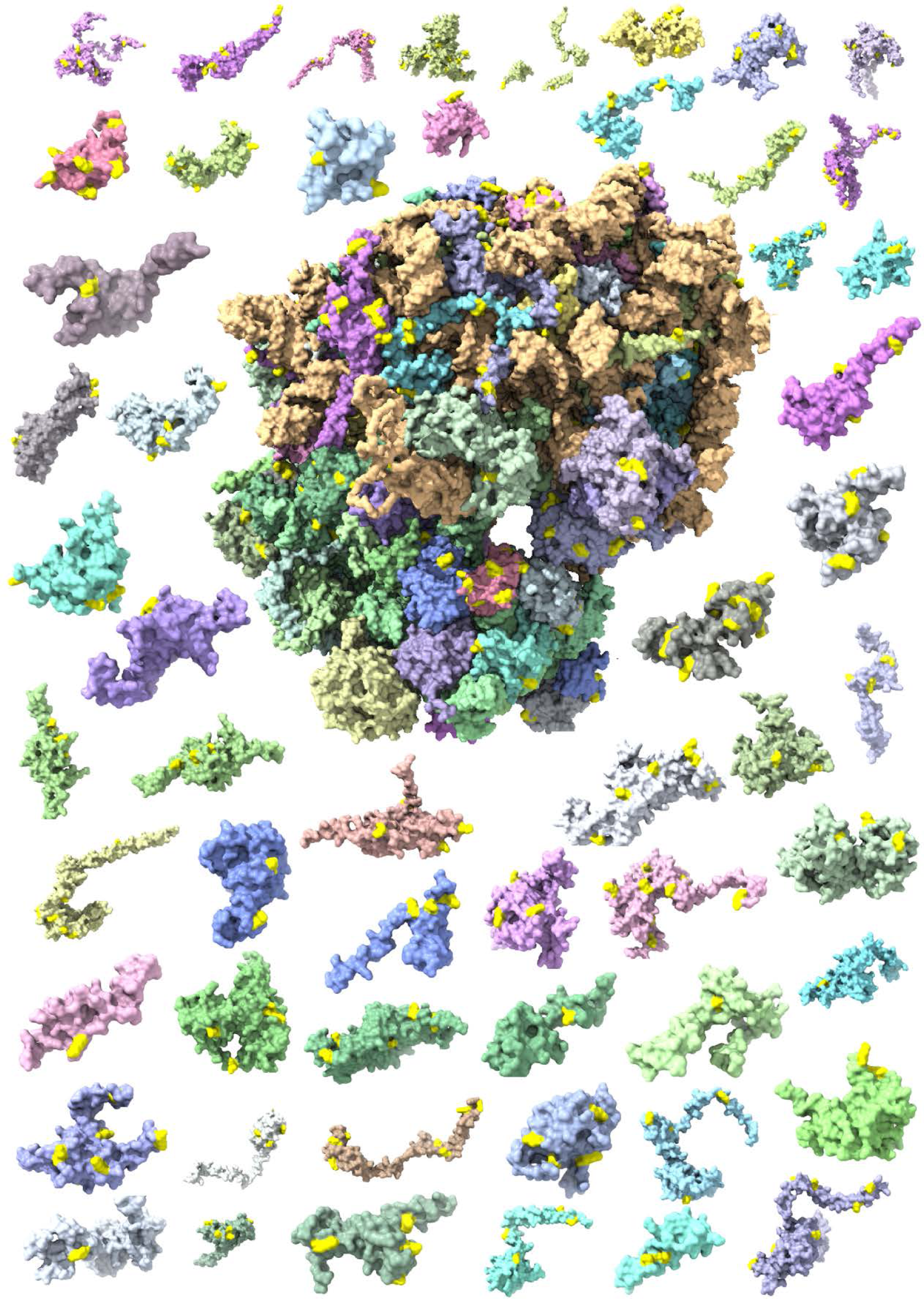
Sites of *In vivo* biotinylations mapped on *in silico* disassembled 80S ribosome (PDB: 6EK0). An “exploded” ribosome plot showing the individual proteins that make up the eukaryotic 80S ribosome with added biotinylation marks (bright yellow) from our 4 biotinylation datasets. We can see that nearly all ribosomal proteins have some yellow “paint” on them.

## Conclusions

We have shown using several orthogonal analyses that *in vivo* biotinylation occurs at a greater rate within predicted IDRs and following on from this observation, highly disordered proteins are more likely to be biotinylated than those that are mostly folded. Furthermore, this trend of increased biotinylation in IDRs is not dependent on the algorithm we use to predict IDRs. However, the greater sensitivity of VSL2b enables the establishment of the trend also in short regions of local disorder and leads to a greater IDR fraction and more biotinylations assigned to IDRs. Finally, we have consistently observed that the SpotBioID study has more proteins that are highly disordered than the other 3 studies thereby validating previous predictions of large fractions of IDRs in nuclear proteins *in vivo*^*48*^.

Moreover, we have interrogated the frequency of post translational modification within IDRs, and have provided an up-to-date analysis of the relationship between *in vivo* observed PTMs across ∼2000 independent experimental studies and predicted IDRs confirming that PTMs are enriched in predicted IDRs. Furthermore, we have shown that the biotinome we have analysed in our study is enriched for PTMs relative to the whole HEK293 proteome despite both groups showing a similar positive correlation between the number of PTMs and number of IDRs. Finally, similar to biotinylation, we find that PTMs too are enriched in nuclear and cytosolic proteins relative to mitochondrial proteins.

## Discussion

We describe here the first *in vivo* evidence for preferential biotinylation of predicted IDRs across four independent proximity proteomics studies. This adds a new type of (exogenous) PTM to a list of other PTMs (phosphorylation, ubiquitination and acetylation) that have previously been suggested to be enriched in IDRs^22,30^ and is validated by our comprehensive analysis. Ubiquitination and acetylation that shares the same target amino acid (i.e. lysine) with most proximity proteomics studies show higher median numbers of modification sites per protein than the deep proteome reference (Fig 3C).

We envisage many possible benefits from re-purposing proximity proteomics data for *in vivo* structure-functional questions:

(i) To complement very detailed kinetic *in vitro* studies that can resolve conformational dynamics at high spatial and temporal resolution using hydrogen deuterium exchange (HDX). Biotin ‘painting’ could enable complementary *in vivo* comparisons of the same target proteins and thereby increase the scope of HDX or related protein surface accessibility-based structural proteomics techniques^54,55^.

(ii) To acquire dynamic snapshots of biological pathways and determine by which mechanism these rewire biomolecular interaction networks and modulate subcellular conformations of proteins. Recent technological advances both in biotinylation enzymes and multiplexed mass analysis will accelerate sampling of more biological timepoints^56-58^.

(iii) To study dynamic *in vivo* drug effects. Many new drug candidates are failing in the later stages of development due to our incomplete understanding of cellular biology. If we can re-purpose BioID or other biotinylation methods for elucidating subcellular protein interactions, we might achieve earlier insights into drug (in)efficiency in relevant biological contexts.

Structural biotin analysis requires identification of the precise sites of biotinylation, which are typically not captured in more widely used protein-level enrichment in BioID experiments. We therefore briefly summarize here possible limitations and benefits for the biotin-peptide enrichment.

An obvious limitation of peptide-level enrichment is that non-biotinylated peptides cannot contribute to the mass spectrometric signal, which can mean that more biological input material may be required in some cases. While peptide-level enrichment increases the specificity and analytical efficiency for detecting biotinylated peptides^36-39^, it comes at the expense of not being able to detect very short proteins that lack lysines or detectable peptides with one missed cleavage (due to a modified lysine). Sequence coverage might be improved by including additional proteases in future biotin-based proximity experiments^59^.

How does biotin painting compare to other recently established proteome-wide structural assays? Similar to (*in vitro/ex vivo*) Limited Proteolysis (LiP)-MS, biotin painting can reveal local structural features of proteins and additionally enables *in vivo* and *in vitro* comparisons while being intrinsically limited by the need for biotin-peptide enrichment^60,61^. (LiP)-MS might, however, be less sensitive for conformational transitions that occur in IDRs that are depleted in hydrophobic amino acids and therefore lack the molecular targets of common LiP enzymes. Thermal proteome profiling (TPP) using quantitative comparisons of soluble fractions upon heating, is also compatible with *in vivo* structural comparisons but lacks local resolution while adding complementary information on protein-protein interactions based on co-precipitation of tightly interacting complex partner proteins at increasing temperatures^21^. Multi-span integral membrane proteins are under-represented in published TPP experiments, and biotin ‘painting’ might have useful complementary applications to biomedically relevant multi-span membrane proteins such as GPCRs. In summary, subcellular biotin painting can complement the already very powerful toolbox of structural proteomics.

Are short IDRs functionally relevant? Several lines of recent independent experimental evidence suggest so. Local flexibility has been identified as crucial factor for the evolution of novel enzymatic functions^62^, and for tuning the activity of enzymes to enable efficient catalysis at low temperatures in biological niches of psychrophilic organisms^63^. Local unfolding, incomplete folding or delayed folding can be helpful for cellular transport of proteins that must not fold prematurely before reaching their cellular destinations^64,65^. High-density biotin painting appears to be useful to characterize the *in vivo* reactivity of both predicted long IDRs (using the IDR predictor IUPred-L) and local or transiently unfolded or disordered IDRs (i.e. IDRs uniquely predicted by VSL2b).

How can we use the insights gleaned from this study to design novel, potentially better, proximity proteomics experiments? A key assumption in classical proximity proteomics studies is that biotinylation is enhanced near the biotin-activating enzyme. Our study shows that unfolded regions can be more readily biotinylated compared to folded regions. This could mean that proteins that in reality never change their cellular distribution can be perceived as farther away or closer to a birA-fusion due to condition-dependent local folding or unfolding, respectively. We do not currently have definitive answers on how to unambiguously dissect condition-dependent local (un)folding and subcellular redistributions. It appears worthwhile to envisage the possibility that transient changes in protein folding can be important modulators of cellular dynamics that should be more broadly factored into experimental designs of proteomics studies (Fig. 6).

**Fig. 6.**
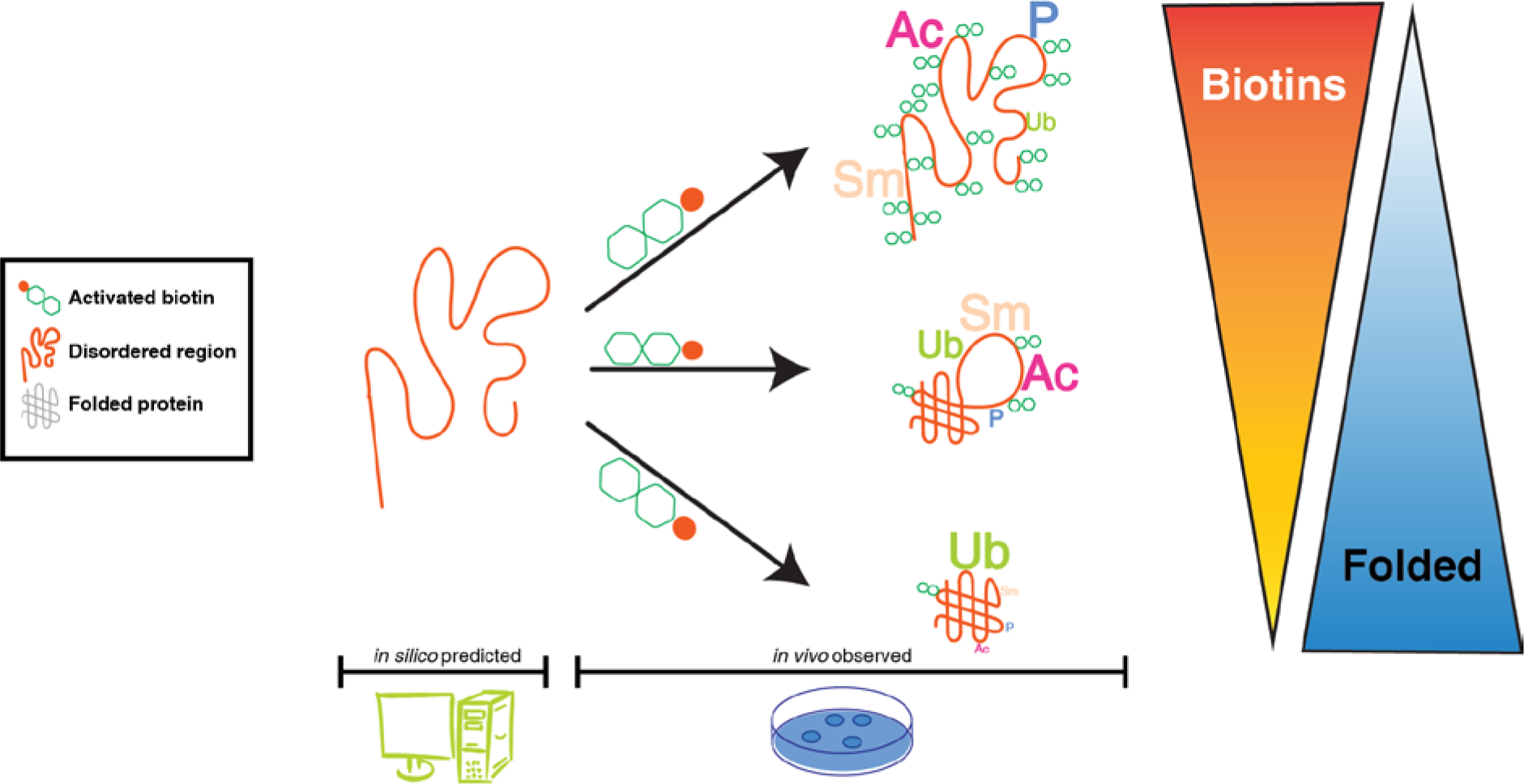
Summarizing model. Biotinylation and other PTMs (including phosphorylation, acetylation, sumoylation, ubiquitination) are more likely in predicted IDRs suggesting that they are more (at least transiently) accessible for biochemical modifications compared to folded proteins. Fully folded proteins can also be modified but show lower fractions of modified residues compared to IDRs except for ubiquitin. This positive correlation of biotinylation density and IDRs, i.e. biotin painting IDRs can be used to re-purpose biotinylation-based proximity proteomics studies to monitor protein plasticity *in vivo*. (clip art modified from: http://www.uidownload.com/free-vectors/fortran-minimalist-monitor-and-computer-clip-art-408587, http://www.clker.com/clipart-red-petri-dish-7.html)

In conclusion, we believe that biotin ‘painting’ adds new layers of insight to proximity proteomics approaches by providing *in vivo* validation for computational IDR predictions, highlighting multi-modification hotspots that are often disease-linked^30^ and by enabling condition and compartment-specific *in vivo* structural comparisons.

## Material and methods

### Source data description

Four independent *in vivo* biotinylation studies have been used for our exploration of structural specificity of biotinylation sites. Their details are provided in Table 1 and they can be accessed as input files on the Github repository https://github.com/ComputationalProteomicsUnit/biotinIDR.

**Table 1:** Sources of data used in this study, Supplementary Information (SI)

| Study | Ref | Target | Chemistry | Data Source | PRIDE ID |
| --- | --- | --- | --- | --- | --- |
| BioSITE | <sup>36</sup> | Tyrosine | APEX2 | SI file 2, supplementary Table 8 | PXD007862 |
| Ab-APEX | <sup>37</sup> | Tyrosine | APEX2 | SI Table 6 | - |
| DiDBiT | <sup>38</sup> | Lysine | NHS-Biotin | SI file 2, Table S27 | - |
| SpotBioID | <sup>39</sup> | Lysine | BirA | SI file 2, sheets 2-5 | - |

### Assigning disorder predictions

Some 60 published disorder prediction algorithms feature balanced accuracies of around 70% to 80%; some being designed and validated to predict short IDRs (<30 residues) and others being better at determining long or both long and short IDRs^66^. The majority of these predictors are trained on a limited set of *in vitro* structural data, mainly X-ray crystallography data, Nuclear Magnetic Resonance (NMR) mobility data in the DisProt database (http://www.disprot.org/)^67^.

A subset of more frequently used prediction algorithms has pre-computed predictions in the web resource D2P2 (http://d2p2.pro/ ^51^). D2P2 also offers the option to select a consensus call for IDRs in a given protein that is predicted by most of the 9 different compound predictors. Of the 9 callers included in D2P2, we focused our interest initially on the 2 most orthogonal callers (1) VSL2b which has high sensitivity for calling IDRs in both short and long regions of IDR^68^, and (2) IUPRed-L which has been trained to predict long disorders with high confidence^52^. As additional comparisons, we also predicted IDRs using (3) a combination of VSL2b and IUPRed-L where an IDR was accepted if called by one or both predictors (4) Consensus of (at least 75% of) 9 predictors included in D2P2.

For all versions of IDR calling, we did not set any restrictions on the length of IDR. This means that an IDR can be called on a residue of length 1. While this might yield a lot of false positives, we wanted to ensure sensitivity rather than specificity of IDR calling. Having tested the 4 different versions of IDR calling with D2P2, we realised consistent trends between all predictions approaches while higher local sensitivity of VSL2b enabled more insights on local disorder. We therefore performed more detailed biotin site and PTM analyses using VSL2b. The IDR assignment uses an Application Programming Interface (API) to the D2P2 website and code to use this API was kindly provided by Dr. Tom Smith. The scripts ‘d2p2.py’ and ‘protinfo.py’ are necessary for the final analysis and can be accessed through the github repository https://github.com/TomSmithCGAT/CamProt/tree/master/camprot/proteomics. The python script for the final IDR analysis and output is called “Get_IDRs-DM-v2.py” and can be accessed via the repository https://github.com/ComputationalProteomicsUnit/biotinIDR.

### Mapping post-Translational Modifications (PTMs)

A full repertoire of PTMs was downloaded on 10^th^ April, 2018 from the “Downloads” section of the PhosphositePlus website (https://www.phosphosite.org/homeAction.action), particularly sites for phosphorylation, acetylation, ubiquitination and sumoylation (Suppl. Tables “PhosphositePlus”). These were then mapped onto the proteins for each of our 4 studies and used for generating protein-wise images.

### Protein sequence modification or proteoform images

Images summarising the location of IDRs and PTMs were produced using Protter (http://wlab.ethz.ch/protter/) and protein structure images were generated using Pymol. For Protter images, scripts were written to generate an appropriate URL and then batch download it from the server. These scripts ‘printUrl.py’ and ‘runUrl.sh’ are also available via the Github repository https://github.com/ComputationalProteomicsUnit/biotinIDR (Suppl. Table “Protter-List”)

### Code availability for Statistical analysis of PTMs and biotinylations in IDRs

Once IDR and PTM information were mapped, data were analysed for correlations and plots were generated using the R statistical framework (https://cran.r-project.org/) and several Bioconductor packages (https://bioconductor.org/). All code and input data can be accessed via the Github respository https://github.com/ComputationalProteomicsUnit/biotinIDR with the main files being ‘biopep.pub.Rmd’ and ‘biopepFunctions.R’. An extension to GO mapping, ‘GO.R’ was also kindly provided by Dr Tom Smith and can be obtained here https://github.com/TomSmithCGAT/CamProt_R.

### Statistical tests used

To compare expected rates of biotin/PTMs and observed counts, we used a standard binomial test in R (binom.test). For estimating the background rate of biotins, we counted all the lysines (K; BirA based studies) or tyrosines (Y; APX based studies) in the protein sequence and within predicted IDR regions. For estimating the background rate of PTMs, we counted all the lysines (K; ubiquitination, acetylation, sumoylation) or serines, threonines and tyrosines (S, T, Y; phosphorylation) in the protein sequence and within predicted IDR regions. We defined the “probability of success” as the number of residues in IDRs/Total number of residues, a “success” as a biotin or PTM within an IDR and “number of trials” is the number of Biotins or PTMs observed in that study.

To look for differences in PTMs and biotins in the three protein groups – Folded, Partially Folded and Unfolded, we used pairwise t-tests or ANOVA followed by a post-hoc correction of family-wise error rates using a Tukey’s Honestly Significant Differences test. The former yields a p-value while the latter yields a confidence interval for the effect size as well as a p-value. To compare number of biotins and IDRs, we used a standard Pearson’s correlation test. To compare mean PTMs between the HEK293 biotinome and HEK293 proteome we used a standard t-test for means.

To perform a GO enrichment analysis, we used the package ‘goseq’^69^ which is based on a Hypergeometric test with a Wallenius’ correction which accounts for any biases in the data such as gene length, protein expression etc. In our study, we used protein expression from Geiger et al. ^53^, as the bias factor prior to calculating GO enrichment.

### Biomolecular structure visualisation

The Cryo-EM structure of the human 80S ribosome (PDB ID 4v6x^70^) was visualised using ChimeraX^71^. SERBP1 and its biotinylated sites were highlighted using the sel function in its command line interface. RNA was coloured purple and protein subunits (except SERBP1) blue. All ribosomal macromolecules were visualised in surface representation. The FKBP3 NMR structure (PDB ID 2mph) was visualised in cartoon model of the first low energy model; surfaces were kept 90% transparent except around biotinylation sites that were highlighted in yellow.

## Code availability

Code to process biotinylation datasets and reproduce the analysis has been deposited in the Github repository https://github.com/ComputationalProteomicsUnit/biotinIDR. Please request access to the code by emailing the corresponding authors as it will be made public following journal acceptance.

## Author Contributions

D.P.M., M.R., K.S.L. conceived the design of the study. D.P.M. and M.R. analysed the data. M.R. mapped and quantified correlations of IDRs, PTMs and sites of biotinylations. D.P.M. inspected sites of biotinylation mapped on experimentally solved high-resolution structures. K.S.L provided insightful feedback along all stages of the project. D.P.M., M.R., K.S.L. wrote the manuscript. D.P.M and M.R contributed equally to the study.

## Acknowledgments

We wish to thank Prof. Keith Dunker and Dr. Sudhakaran Prabakaran for inspiring discussions, advice and comments on earlier versions of this manuscript. We thank Dewi Eburne for help with experiments that triggered our interest in structural features of biotinomes, Dr. Tom Smith for kindly sharing his code for IDR parsing and GO term extension, and Dr. Eneko Villanueva and Dr. Rayner Queiroz and Prof. Ben Luisi for their insightful discussions. We thank Dr. Matthew Young for contributing his ideas on complementary data analysis strategies. D.P.M. is supported by the BBSRC grant BB/N010493/1 and a TMT grant (Thermo Fisher, 2018) for studying cellular biotinylations.

## Additional information

### Supplementary Information

All supplementary tables are available as Excel sheets. A supplementary video (mp4) is available as well.

### Competing interests

The authors have declared no conflict of interest.

**Fig. S1.**
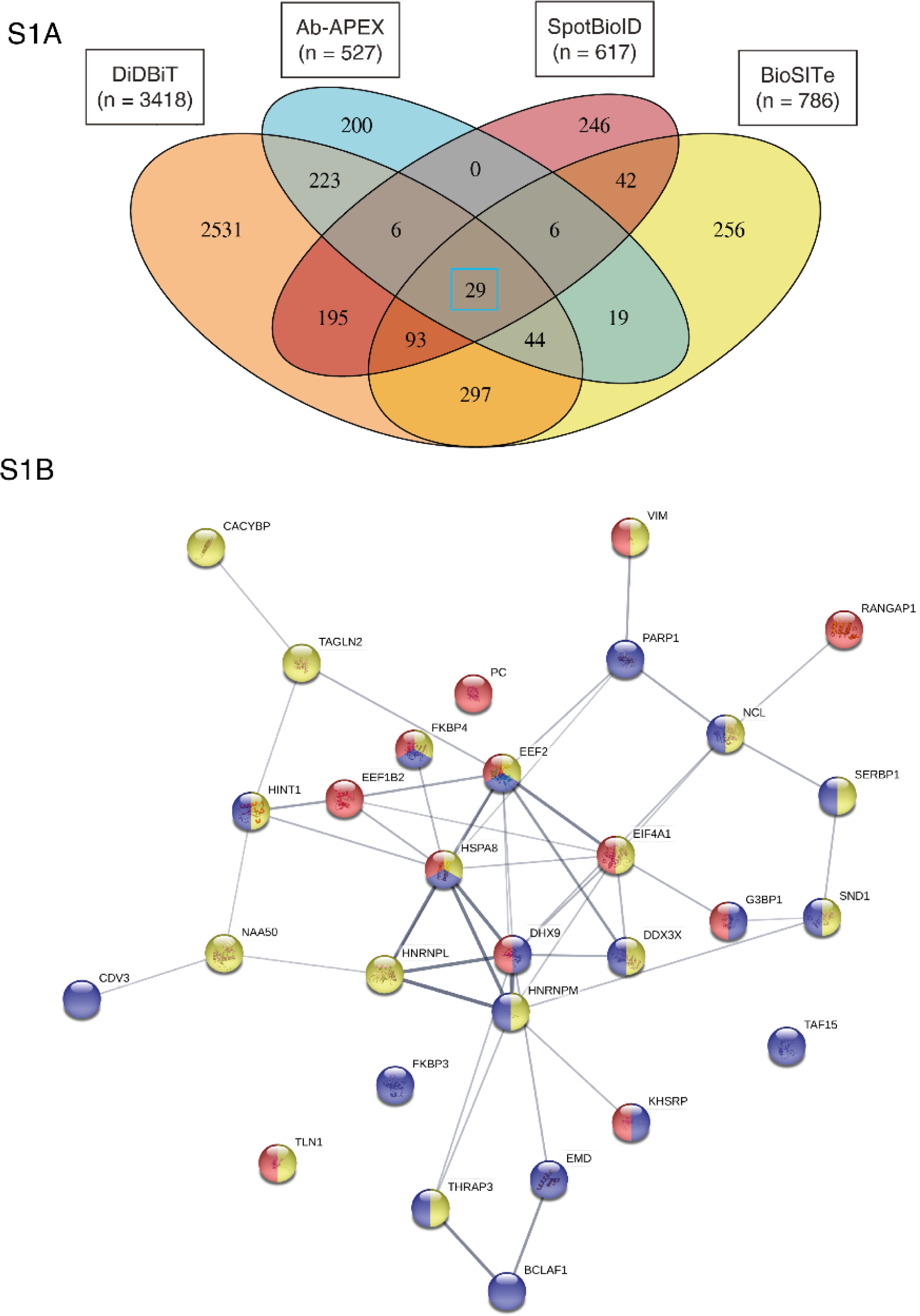
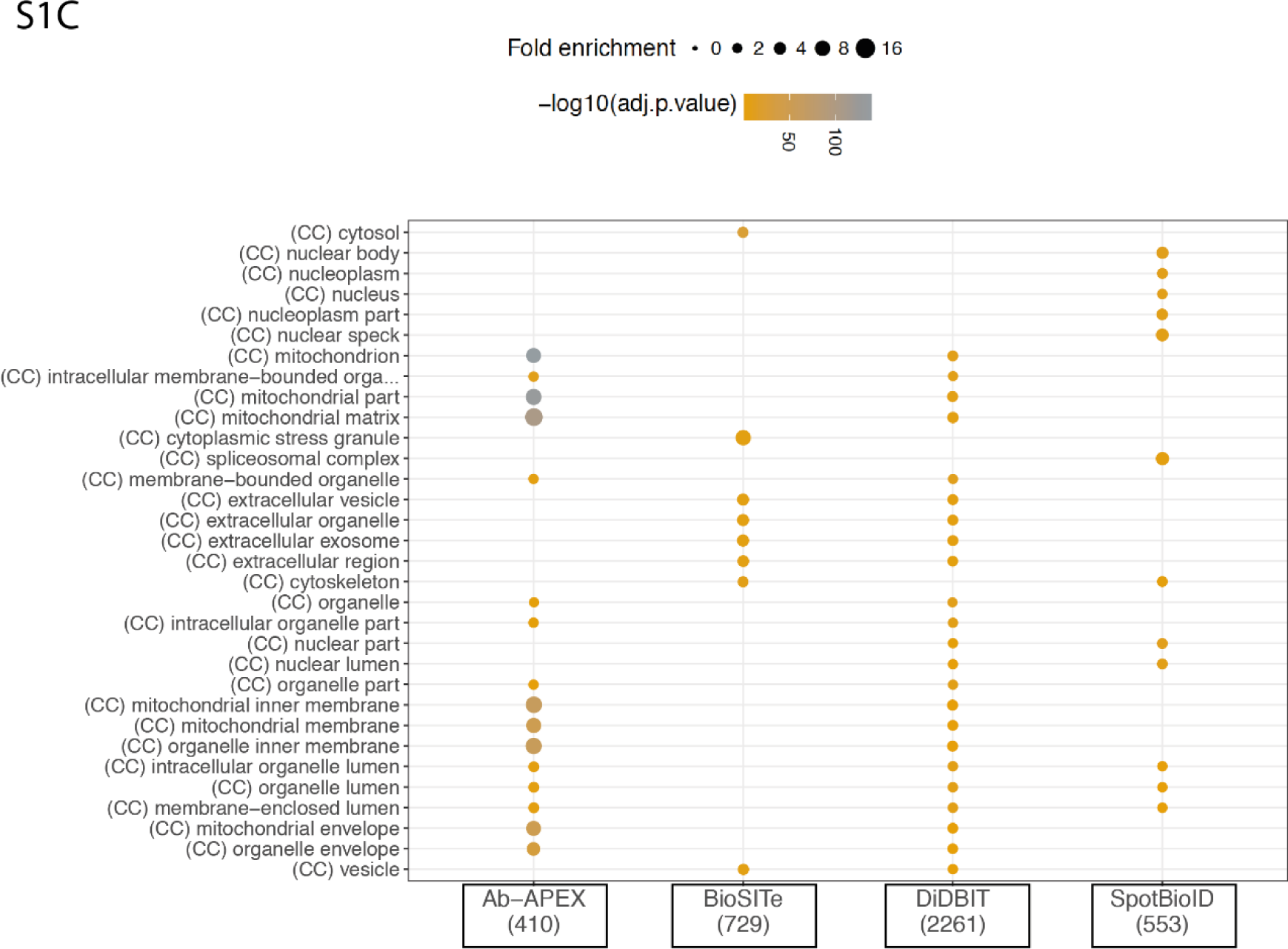
**(A)** A Venn diagram showing the overlap of proteins across the four studies used in this analysis. There is a very small number of proteins (29) common to all 4 studies (blue box). DiDBIT has the most number of exclusive proteins as it targets the entire cell. Ab-APEX targets the mitochondrial matrix and inner mitochondrial membrane. SpotBioID targets the nucleus and BioSITE targets the cytoplasm. **(B)** A connectivity plot showing any published and validated interactions between the 29 common proteins identified in S1A. This image was generated using the online program STRING (https://string-db.org/cgi/input.pl?sessionId=QqkSTNv1EVki&input_page_show_search=on). An enrichment analysis was run on the 29 proteins using Gene Ontology (GO) categories and the colours represent some of the most significant terms from this analysis. Blue indicates that these proteins are known to localise to the nucleus, red indicates localisation to the cytosol and yellow indicates localisation to the extracellular region. Multiple colours in a single circle indicate that the given protein has been found in multiple locations in different studies. **(C)** GO Cellular Component enrichment analysis for the 4 studies. The proteins in each study were mapped to GO:CC categories and compared to a background of the published HEK293 proteome which was also mapped to GO:CC catagories. The size of the dot represents the fold enrichment over the background, i.e. the fraction of proteins in the input list that are annotated by a given GO term divided by fraction of proteins in the background list that could be mapped to the same GO term. The colour of the dot represents the significance of the enrichment with grey being most and orange being least significant. Note that all terms displayed in this figure are significant and above the adjusted p-value cut-off of 0.05.

**Fig. S2.**
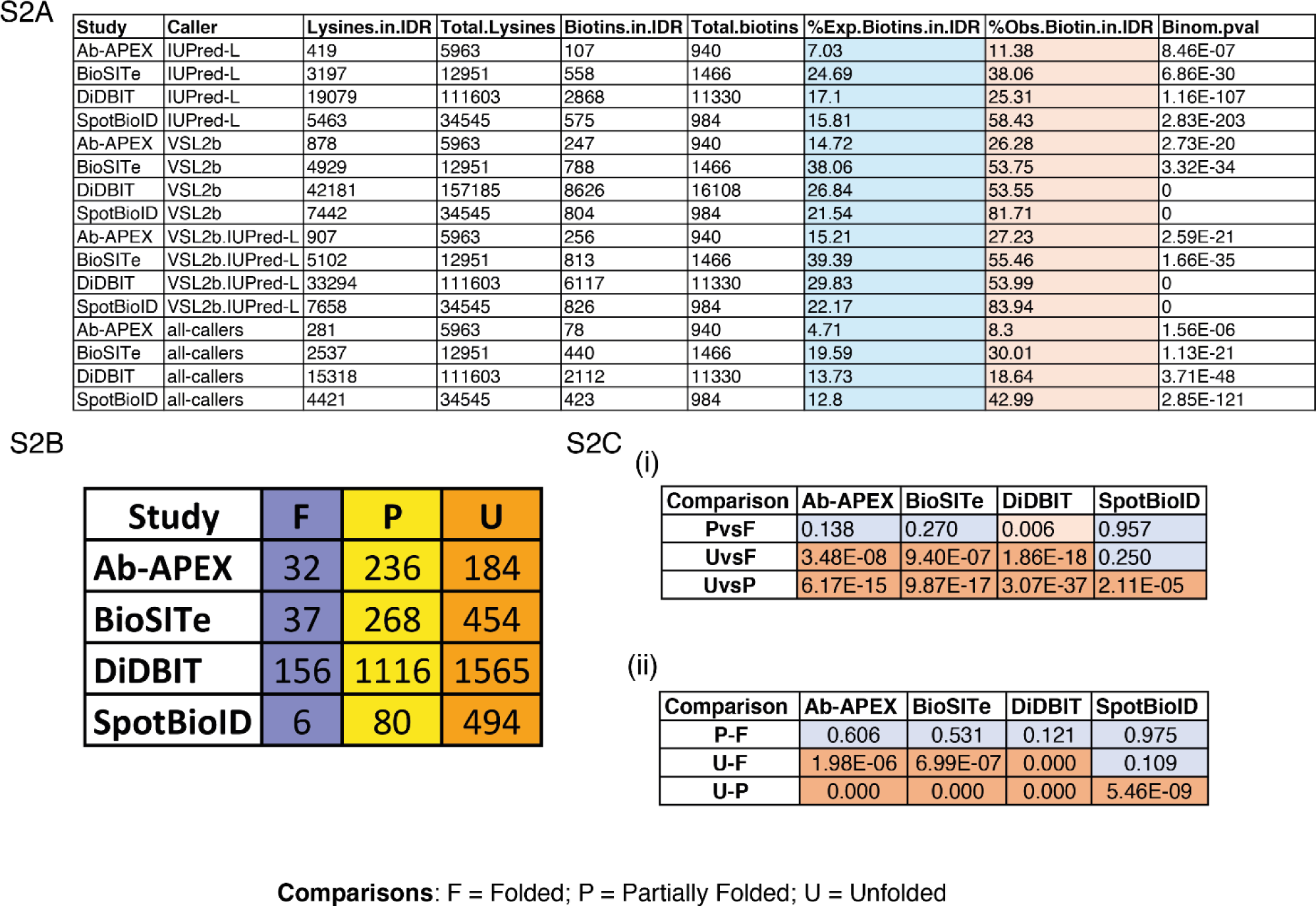
Statistics for Biotin and IDR comparisons. **(A)** A table of values and percentages used to look at the Expected (Blue) and Observed (Orange) rate of biotin occurrence within IDRs across all studies and all callers. The last column “Binom.pval” denotes the p-value using a bionomial test where the “probability of success” is the (Lysine residues in IDRs/Total Lysine residues), a “success” is a biotin within an IDR (Biotins.in.IDR) and “number of trials” is the total number of biotins (Total.biotins) observed in that study. All tests are significant at the p = 0.05 level **(B)** A table displaying the frequencies of proteins in each of the IDR categories in each of the 4 studies using the IDR predictor VSL2b. **(C)** To test whether or not there were significant differences in the number of biotins found in regions of IDR across the 3 IDR categories **(i)** a pair-wise t-test with multiple testing correction between the three groups – F, P and U. The table shows the p-value of these pairwise t-tests across the four studies. Light blue denotes comparisons that are not significant. Light orange denotes significant (p<0.05) and dark orange denotes comparisons that are highly significant (p << 0.05) **(ii)** an analysis of variance (ANOVA) was performed followed by a Tukey Honestly Significant Differences (THSD) test to correct for family wise error. The table shows the p-value of the THSD test across the four studies. Light blue denotes comparisons that are not significant. Light orange denotes significant (p<0.05) and dark orange denotes comparisons that are highly significant (p << 0.05)

**Fig. S3.**
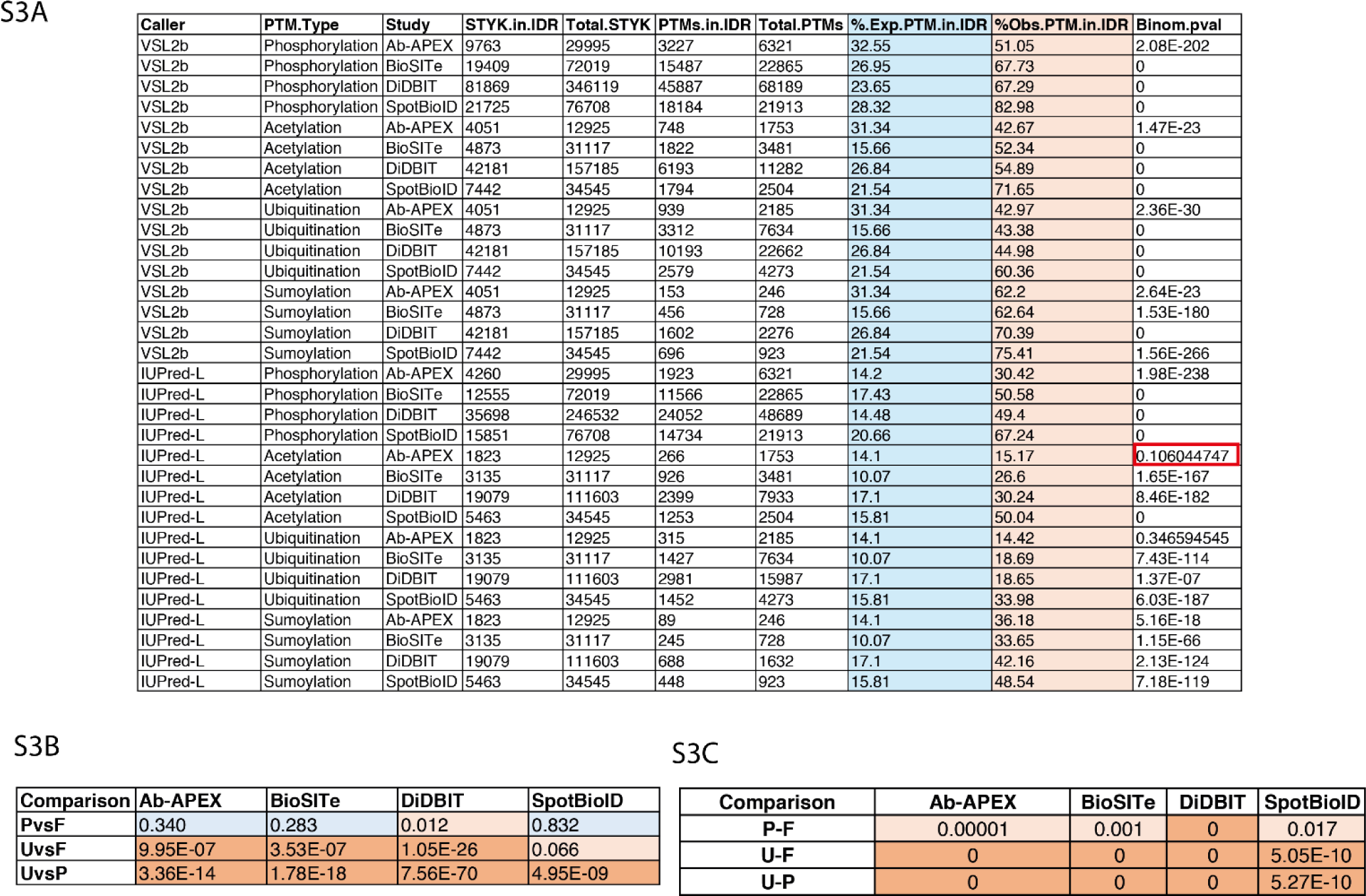
Statistics for PT and IDR comparisons. (**A)** A table of values and percentages used to look at the Expected (Blue) and Observed (Orange) rate of PTM occurrence within IDRs across all studies, all callers and all PTM types. The “Binom.pval” column denotes p-values obtained using a bionomial test where the “probability of success” is the (STKY.in.IDR/Total.STKY), a “success” is a PTM within an IDR (PTMS.in.IDRs) and “number of trials” is the number of (each type of) PTMs (Total.PTMs) observed in that study (Fig. S3A)**. (B)** a pair-wise t-test with multiple testing correction between the three groups – F, P and U. The table shows the p-value of these pairwise t-tests across the four studies. Light blue denotes comparisons that are not significant. Light orange denotes significant (p<0.05) and dark orange denotes comparisons that are highly significant (p << 0.05) **(C)** an analysis of variance (ANOVA) followed by a Tukey Honestly Significant Differences (THSD) test to correct for family wise error. The table shows the p-value of the THSD test across the four studies. Light blue denotes comparisons that are not significant. Light orange denotes significant (p<0.05) and dark orange denotes comparisons that are highly significant (p << 0.05).

## References

1 Yruela, I., Oldfield, C. J., Niklas, K. J. & Dunker, A. K. Evidence for a Strong Correlation Between Transcription Factor Protein Disorder and Organismic Complexity. Genome Biol Evol 9, 1248–1265, doi:10.1093/gbe/evx073 (2017).

2 Xue, B., Dunker, A. K. & Uversky, V. N. Orderly order in protein intrinsic disorder distribution: disorder in 3500 proteomes from viruses and the three domains of life. J Biomol Struct Dyn 30, 137–149, doi:10.1080/07391102.2012.675145 (2012).

3 Romero, P. R. et al. Alternative splicing in concert with protein intrinsic disorder enables increased functional diversity in multicellular organisms. Proc Natl Acad Sci U S A 103, 8390–8395, doi:10.1073/pnas.0507916103 (2006).

4 Dunker, A. K., Bondos, S. E., Huang, F. & Oldfield, C. J. Intrinsically disordered proteins and multicellular organisms. Semin Cell Dev Biol 37, 44–55, doi:10.1016/j.semcdb.2014.09.025 (2015).

5 Minde, D. P., Dunker, A. K. & Lilley, K. S. Time, space and disorder in the expanding proteome universe. Proteomics, doi:10.1002/pmic.201600399 (2017).

6 Tompa, P., Davey, N. E., Gibson, T. J. & Babu, M. M. A million peptide motifs for the molecular biologist. Mol Cell 55, 161–169, doi:10.1016/j.molcel.2014.05.032 (2014).

7 Smith, L. M., Kelleher, N. L. & Consortium for Top Down, P. Proteoform: a single term describing protein complexity. Nat Methods 10, 186–187, doi:10.1038/nmeth.2369 (2013).

8 Uversky, V. N., Oldfield, C. J. & Dunker, A. K. Intrinsically disordered proteins in human diseases: introducing the D2 concept. Annu Rev Biophys 37, 215–246, doi:10.1146/annurev.biophys.37.032807.125924 (2008).

9 Minde, D. P., Anvarian, Z., Rudiger, S. G. D. & Maurice, M. M. Messing up disorder: how do missense mutations in the tumor suppressor protein APC lead to cancer? Molecular Cancer 10, doi:10.1186/1476-4598-10-101 (2011).

10 Noutsou, M. et al. Critical Scaffolding Regions of the Tumor Suppressor Axin1 Are Natively Unfolded. Journal of Molecular Biology 405, 773–786, doi:10.1016/j.jmb.2010.11.013 (2011).

11 Minde, D. P., Radli, M., Forneris, F., Maurice, M. M. & Ruediger, S. G. D. Large Extent of Disorder in Adenomatous Polyposis Coli Offers a Strategy to Guard Wnt Signalling against Point Mutations. Plos One 8, doi:10.1371/journal.pone.0077257 (2013).

12 Carulla, N. et al. Experimental characterization of disordered and ordered aggregates populated during the process of amyloid fibril formation. Proc Natl Acad Sci U S A 106, 7828–7833, doi:10.1073/pnas.0812227106 (2009).

13 Folkers, P. J. et al. Solution structure of recombinant hirudin and the Lys-47 Glu mutant: a nuclear magnetic resonance and hybrid distance geometry-dynamical simulated annealing study. Biochemistry 28, 2601–2617 (1989).

14 Vucetic, S., Brown, C. J., Dunker, A. K. & Obradovic, Z. Flavors of protein disorder. Proteins 52, 573–584, doi:10.1002/prot.10437 (2003).

15 Babu, M. M. The contribution of intrinsically disordered regions to protein function, cellular complexity, and human disease. Biochem Soc Trans 44, 1185–1200, doi:10.1042/BST20160172 (2016).

16 Toth-Petroczy, A. et al. Structured States of Disordered Proteins from Genomic Sequences. Cell 167, 158–170 e112, doi:10.1016/j.cell.2016.09.010 (2016).

17 Gunasekaran, K., Tsai, C. J. & Nussinov, R. Analysis of ordered and disordered protein complexes reveals structural features discriminating between stable and unstable monomers. J Mol Biol 341, 1327–1341, doi:10.1016/j.jmb.2004.07.002 (2004).

18 Gsponer, J., Futschik, M. E., Teichmann, S. A. & Babu, M. M. Tight regulation of unstructured proteins: from transcript synthesis to protein degradation. Science 322, 1365–1368, doi:10.1126/science.1163581 (2008).

19 Dyson, H. J. & Wright, P. E. Intrinsically unstructured proteins and their functions. Nat Rev Mol Cell Biol 6, 197–208, doi:10.1038/nrm1589 (2005).

20 Borgia, A. et al. Extreme disorder in an ultrahigh-affinity protein complex. Nature 555, 61–66, doi:10.1038/nature25762 (2018).

21 Becher, I. et al. Pervasive Protein Thermal Stability Variation during the Cell Cycle. Cell 173, 1495–1507 e1418, doi:10.1016/j.cell.2018.03.053 (2018).

22 Iakoucheva, L. M. et al. The importance of intrinsic disorder for protein phosphorylation. Nucleic Acids Res 32, 1037–1049, doi:10.1093/nar/gkh253 (2004).

23 Zhu, S. et al. Hyperphosphorylation of intrinsically disordered tau protein induces an amyloidogenic shift in its conformational ensemble. PLoS One 10, e0120416, doi:10.1371/journal.pone.0120416 (2015).

24 Uversky, V. N. The intrinsic disorder alphabet. III. Dual personality of serine. Intrinsically Disord Proteins 3, e1027032, doi:10.1080/21690707.2015.1027032 (2015).

25 Kulkarni, P. et al. Phosphorylation-induced conformational dynamics in an intrinsically disordered protein and potential role in phenotypic heterogeneity. Proc Natl Acad Sci U S A 114, E2644–E2653, doi:10.1073/pnas.1700082114 (2017).

26 Potel, C. M., Lin, M. H., Heck, A. J. R. & Lemeer, S. Widespread bacterial protein histidine phosphorylation revealed by mass spectrometry-based proteomics. Nat Methods 15, 187–190, doi:10.1038/nmeth.4580 (2018).

27 Rosenlow, J., Isaksson, L., Mayzel, M., Lengqvist, J. & Orekhov, V. Y. Tyrosine phosphorylation within the intrinsically disordered cytosolic domains of the B-cell receptor: an NMR-based structural analysis. PLoS One 9, e96199, doi:10.1371/journal.pone.0096199 (2014).

28 Guharoy, M., Bhowmick, P. & Tompa, P. Design Principles Involving Protein Disorder Facilitate Specific Substrate Selection and Degradation by the Ubiquitin-Proteasome System. J Biol Chem 291, 6723–6731, doi:10.1074/jbc.R115.692665 (2016).

29 Kim, S. C. et al. Substrate and functional diversity of lysine acetylation revealed by a proteomics survey. Mol Cell 23, 607–618, doi:10.1016/j.molcel.2006.06.026 (2006).

30 Huang, Q. et al. Human proteins with target sites of multiple post-translational modification types are more prone to be involved in disease. J Proteome Res 13, 2735–2748, doi:10.1021/pr401019d (2014).

31 Han, S. et al. Proximity Biotinylation as a Method for Mapping Proteins Associated with mtDNA in Living Cells. Cell Chem Biol 24, 404–414, doi:10.1016/j.chembiol.2017.02.002 (2017).

32 Rees, J. S., Li, X. W., Perrett, S., Lilley, K. S. & Jackson, A. P. Protein Neighbors and Proximity Proteomics. Mol Cell Proteomics 14, 2848–2856, doi:10.1074/mcp.R115.052902 (2015).

33 Kim, D. I. et al. An improved smaller biotin ligase for BioID proximity labeling. Mol Biol Cell 27, 1188–1196, doi:10.1091/mbc.E15-12-0844 (2016).

34 Tron, C. M. et al. Structural and functional studies of the biotin protein ligase from Aquifex aeolicus reveal a critical role for a conserved residue in target specificity. J Mol Biol 387, 129–146 (2009).

35 Youn, J. Y. et al. High-Density Proximity Mapping Reveals the Subcellular Organization of mRNA-Associated Granules and Bodies. Mol Cell 69, 517–532 e511, doi:10.1016/j.molcel.2017.12.020 (2018).

36 Kim, D. I. et al. BioSITe: A Method for Direct Detection and Quantitation of Site-Specific Biotinylation. J Proteome Res 17, 759–769, doi:10.1021/acs.jproteome.7b00775 (2018).

37 Udeshi, N. D. et al. Antibodies to biotin enable large-scale detection of biotinylation sites on proteins. Nat Methods 14, 1167–1170, doi:10.1038/nmeth.4465 (2017).

38 Schiapparelli, L. M. et al. Direct detection of biotinylated proteins by mass spectrometry. J Proteome Res 13, 3966–3978, doi:10.1021/pr5002862 (2014).

39 Lee, S. Y. et al. Proximity-Directed Labeling Reveals a New Rapamycin-Induced Heterodimer of FKBP25 and FRB in Live Cells. ACS Cent Sci 2, 506–516, doi:10.1021/acscentsci.6b00137 (2016).

40 Lam, S. S. et al. Directed evolution of APEX2 for electron microscopy and proximity labeling. Nat Methods 12, 51–54, doi:10.1038/nmeth.3179 (2015).

41 Roux, K. J., Kim, D. I. & Burke, B. BioID: a screen for protein-protein interactions. Curr Protoc Protein Sci 74, Unit 19 23, doi:10.1002/0471140864.ps1923s74 (2013).

42 Campen, A. et al. TOP-IDP-scale: a new amino acid scale measuring propensity for intrinsic disorder. Protein Pept Lett 15, 956–963 (2008).

43 Sugase, K., Dyson, H. J. & Wright, P. E. Mechanism of coupled folding and binding of an intrinsically disordered protein. Nature 447, 1021, doi:10.1038/nature05858 (2007).

44 Davey, N. E. et al. Attributes of short linear motifs. Mol Biosyst 8, 268–281, doi:10.1039/c1mb05231d (2012).

45 Miskei, M., Antal, C. & Fuxreiter, M. FuzDB: database of fuzzy complexes, a tool to develop stochastic structure-function relationships for protein complexes and higher-order assemblies. Nucleic Acids Res 45, D228–D235, doi:10.1093/nar/gkw1019 (2017).

46 Monteiro, R. et al. Differential biotin labelling of the cell envelope proteins in lipopolysaccharidic diderm bacteria: Exploring the proteosurfaceome of Escherichia coli using sulfo-NHS-SS-biotin and sulfo-NHS-PEG4-bismannose-SS-biotin. J Proteomics, doi:10.1016/j.jprot.2018.03.026 (2018).

47 Lins, L., Thomas, A. & Brasseur, R. Analysis of accessible surface of residues in proteins. Protein Sci 12, 1406–1417, doi:10.1110/ps.0304803 (2003).

48 Frege, T. & Uversky, V. N. Intrinsically disordered proteins in the nucleus of human cells. Biochem Biophys Rep 1, 33–51, doi:10.1016/j.bbrep.2015.03.003 (2015).

49 Peng, Z. et al. A creature with a hundred waggly tails: intrinsically disordered proteins in the ribosome. Cell Mol Life Sci 71, 1477–1504, doi:10.1007/s00018-013-1446-6 (2014).

50 Ito, M. et al. Intrinsically disordered proteins in human mitochondria. Genes Cells 17, 817–825, doi:10.1111/gtc.12000 (2012).

51 Oates, M. E. et al. D(2)P(2): database of disordered protein predictions. Nucleic Acids Res 41, D508–516, doi:10.1093/nar/gks1226 (2013).

52 Dosztanyi, Z., Csizmok, V., Tompa, P. & Simon, I. IUPred: web server for the prediction of intrinsically unstructured regions of proteins based on estimated energy content. Bioinformatics 21, 3433–3434, doi:10.1093/bioinformatics/bti541 (2005).

53 Geiger, T., Wehner, A., Schaab, C., Cox, J. & Mann, M. Comparative proteomic analysis of eleven common cell lines reveals ubiquitous but varying expression of most proteins. Mol Cell Proteomics 11, M111 014050, doi:10.1074/mcp.M111.014050 (2012).

54 Gustavsson, M. et al. Structural basis of ligand interaction with atypical chemokine receptor 3. Nat Commun 8, 14135, doi:10.1038/ncomms14135 (2017).

55 Gladkova, C., Maslen, S., Skehel, J. M. & Komander, D. Mechanism of parkin activation by PINK1. Nature, doi:10.1038/s41586-018-0224-x (2018).

56 Ramanathan, M. et al. RNA–protein interaction detection in living cells. Nature Methods 15, 207, doi:10.1038/nmeth.4601 (2018).

57 Virreira Winter, S. et al. EASI-tag enables accurate multiplexed and interference-free MS2-based proteome quantification. Nat Methods, doi:10.1038/s41592-018-0037-8 (2018).

58 Kelstrup, C. D. et al. Performance Evaluation of the Q Exactive HF-X for Shotgun Proteomics. J Proteome Res 17, 727–738, doi:10.1021/acs.jproteome.7b00602 (2018).

59 Meyer, J. G. et al. Expanding proteome coverage with orthogonal-specificity alpha-lytic proteases. Mol Cell Proteomics 13, 823–835, doi:10.1074/mcp.M113.034710 (2014).

60 Feng, Y. et al. Global analysis of protein structural changes in complex proteomes. Nature Biotechnology 32, 1036, doi:10.1038/nbt.2999 (2014).

61 Piazza, I. et al. A Map of Protein-Metabolite Interactions Reveals Principles of Chemical Communication. Cell 172, 358–372 e323, doi:10.1016/j.cell.2017.12.006 (2018).

62 Campbell, E. et al. The role of protein dynamics in the evolution of new enzyme function. Nature Chemical Biology 12, 944, doi:10.1038/nchembio.2175 (2016).

63 Saavedra, H. G., Wrabl, J. O., Anderson, J. A., Li, J. & Hilser, V. J. Dynamic allostery can drive cold adaptation in enzymes. Nature 558, 324–328, doi:10.1038/s41586-018-0183-2 (2018).

64 Zhou, J. & Dunker, A. K. Regulating Protein Function by Delayed Folding. Structure 26, 679–681, doi:10.1016/j.str.2018.04.011 (2018).

65 Tsirigotaki, A. et al. Long-Lived folding intermediates predominate the Targeting-Competent secretome. Structure 26, 695-707. e695 (2018).

66 Peng, Z. L. & Kurgan, L. Comprehensive comparative assessment of in-silico predictors of disordered regions. Curr Protein Pept Sci 13, 6–18 (2012).

67 Piovesan, D. et al. DisProt 7.0: a major update of the database of disordered proteins. Nucleic Acids Research, gkw1056 (2016).

68 Peng, K., Radivojac, P., Vucetic, S., Dunker, A. K. & Obradovic, Z. Length-dependent prediction of protein intrinsic disorder. BMC Bioinformatics 7, 208, doi:10.1186/1471-2105-7-208 (2006).

69 Young, M. D., Wakefield, M. J., Smyth, G. K. & Oshlack, A. Gene ontology analysis for RNA-seq: accounting for selection bias. Genome Biology 11, R14, doi:10.1186/gb-2010-11-2-r14 (2010).

70 Anger, A. M. et al. Structures of the human and Drosophila 80S ribosome. Nature 497, 80–85, doi:10.1038/nature12104 (2013).

71 Goddard, T. D. et al. UCSF ChimeraX: Meeting modern challenges in visualization and analysis. Protein Sci 27, 14–25, doi:10.1002/pro.3235 (2018).

72 Kampstra, P. Beanplot: A Boxplot Alternative for Visual Comparison of Distributions. 2008 28, 9, doi:10.18637/jss.v028.c01 (2008).

